# Cloning of the rice *Xo1* resistance gene and interaction of the Xo1 protein with the defense-suppressing *Xanthomonas* effector Tal2h

**DOI:** 10.1101/2020.05.26.116731

**Authors:** Andrew C. Read, Mathilde Hutin, Matthew J. Moscou, Fabio C. Rinaldi, Adam J. Bogdanove

**Affiliations:** Plant Pathology and Plant-Microbe Biology Section, School of Integrative Plant Science, Cornell University, Ithaca, NY 14853; IRD, CIRAD, Université Montpellier, IPME, 34000 Montpellier, France; The Sainsbury Laboratory, University of East Anglia, Norwich Research Park, Norwich, NR4 7UK, United Kingdom

**Keywords:** Resistance genes, effectors, defense suppression, nucleotide binding leucine-rich repeat (NLR), transcription activator-like effector (TALE), truncTALE, mass spectrometry, protein-protein interaction

## Abstract

The *Xo1* locus in the heirloom rice variety Carolina Gold Select confers resistance to bacterial leaf streak and bacterial blight, caused by *Xanthomonas oryzae* pvs. oryzicola and oryzae, respectively. Resistance is triggered by pathogen-delivered transcription activator-like effectors (TALEs) independent of their ability to activate transcription, and is suppressed by variants called truncTALEs common among Asian strains. By transformation of the susceptible variety Nipponbare, we show that one of 14 nucleotide-binding, leucine-rich repeat (NLR) protein genes at the locus, with a zfBED domain, is the *Xo1* gene. Analyses of published transcriptomes revealed that the *Xo1*-mediated response is similar to those of NLR resistance genes *Pia* and *Rxo1* and distinct from that associated with induction of the executor resistance gene *Xa23*, and that a truncTALE dampens or abolishes activation of defense-associated genes by *Xo1*. In *Nicotiana benthamiana* leaves, fluorescently-tagged Xo1 protein, like TALEs and truncTALEs, localized to the nucleus. And, endogenous Xo1 specifically co-immunoprecipitated from rice leaves with a pathogen-delivered, epitope-tagged truncTALE. These observations suggest that suppression of Xo1-function by truncTALEs occurs through direct or indirect physical interaction. They further suggest that effector co-immunoprecipitation may be effective for identifying or characterizing other resistance genes.

---

Bacterial leaf streak of rice, caused by *Xanthomonas oryzae* pv. oryzicola (Xoc), is an increasing threat to production in many parts of the world, especially in Africa. Bacterial blight of rice, caused by *X. oryzae* pv. oryzae (Xoo) has long been a major constraint in Asia and is becoming prevalent in Africa. The purified American heirloom rice variety Carolina Gold Select (hereafter Carolina Gold; McClung and Fjellstrom, 2010) is resistant to all tested African strains of Xoc and some tested strains of Xoo (Read et al., 2016). Using an African strain of Xoc, the resistance was mapped to chromosome 4 and designated as *Xo1* (Triplett et al., 2016). Both Xoc and Xoo deploy multiple type III-secreted transcription activator-like effectors (TALEs) during infection. TALEs enter the plant nucleus and bind to promoters, each with different sequence specificity, to transcriptionally activate effector-specific target genes (Perez-Quintero and Szurek, 2019). Some of these genes, called susceptibility genes, contribute to disease development (Hutin et al., 2015). In some host genotypes, a TALE may activate a so-called executor resistance gene, leading to host cell death that stops the infection (Bogdanove et al., 2010). Most of the cloned resistance genes for bacterial blight are in fact executor genes (Zhang et al., 2015). *Xo1* is different. It mediates resistance in response to TALEs with distinct DNA-binding specificities independent of their ability to activate transcription (Triplett et al., 2016). Also, unlike executor genes, *Xo1* function is suppressed by a variant class of these effectors known as truncTALEs (also called iTALEs). Like TALEs, TruncTALEs nuclear localize (Ji et al., 2016), however due to large N and C terminal deletions they do not bind DNA (Read et al., 2016).

*Xo1* maps to a region that in the reference rice genome (cv. Nipponbare) contains seven nucleotide-binding, leucine-rich repeat protein genes (“NLR” genes) (Triplett et al., 2016). NLR genes are the largest class of plant disease resistance genes. NLR proteins recognize specific, corresponding pathogen effector proteins through direct interaction or by detecting effector-dependent changes of host target proteins, and mediate downstream defense signaling that leads to expression of defense genes and a programmed localized cell death, the hypersensitive reaction (HR) (Lolle et al., 2020). Recently, by whole genome sequencing, we determined that the *Xo1* locus in Carolina Gold comprises 14 NLR genes. We identified one of these, *Xo1_11_*, as a strong candidate based on its structural similarity to the previously cloned and only known NLR resistance gene for bacterial blight, *Xa1* (Read et al., 2020). *Xa1*, originally identified in the rice variety Kogyoku, maps to the same location (Yoshimura et al., 1998) and behaves similarly to *Xo1*: it mediates recognition of TALEs with distinct DNA-binding specificities (and thus confers resistance also to bacterial leaf streak), and its activity is suppressed by truncTALEs (Ji et al., 2016). *Xo1_11_* and *Xa1* are members of a small subfamily of NLR genes that encode an unusual N-terminal domain comprising a zinc finger BED (zfBED) motif (Read et al., 2020).

To ascertain whether *Xo1_11_* is the gene responsible for *Xo1* resistance, we generated transgenic Nipponbare plants expressing it. For transformation, we amplified the genomic *Xo1_11_* coding sequence (5,882 bp) as well as the 993 bp region upstream of the start codon and cloned them together into a binary vector with a 35S terminator. T0 *Xo1_11_* plants were inoculated by syringe infiltration with African Xoc strain CFBP7331, which has no truncTALE of its own, carrying either an empty vector (EV) or the plasmid-borne truncTALE gene *tal2h* (p2h) from the Asian Xoc strain BLS256 (Read et al., 2016). Phenotypes of CFBP7331(EV) and CFBP7331(p2h) were confirmed on untransformed Nipponbare and Carolina Gold plants (**Fig. S1**). Plants from two *Xo1_11_* transformation events displayed resistance to the strain with the EV, but not to the strain carrying Tal2h (**Fig. 1**), demonstrating that *Xo1_11_* is the *Xo1* gene.

**Fig. 1.**
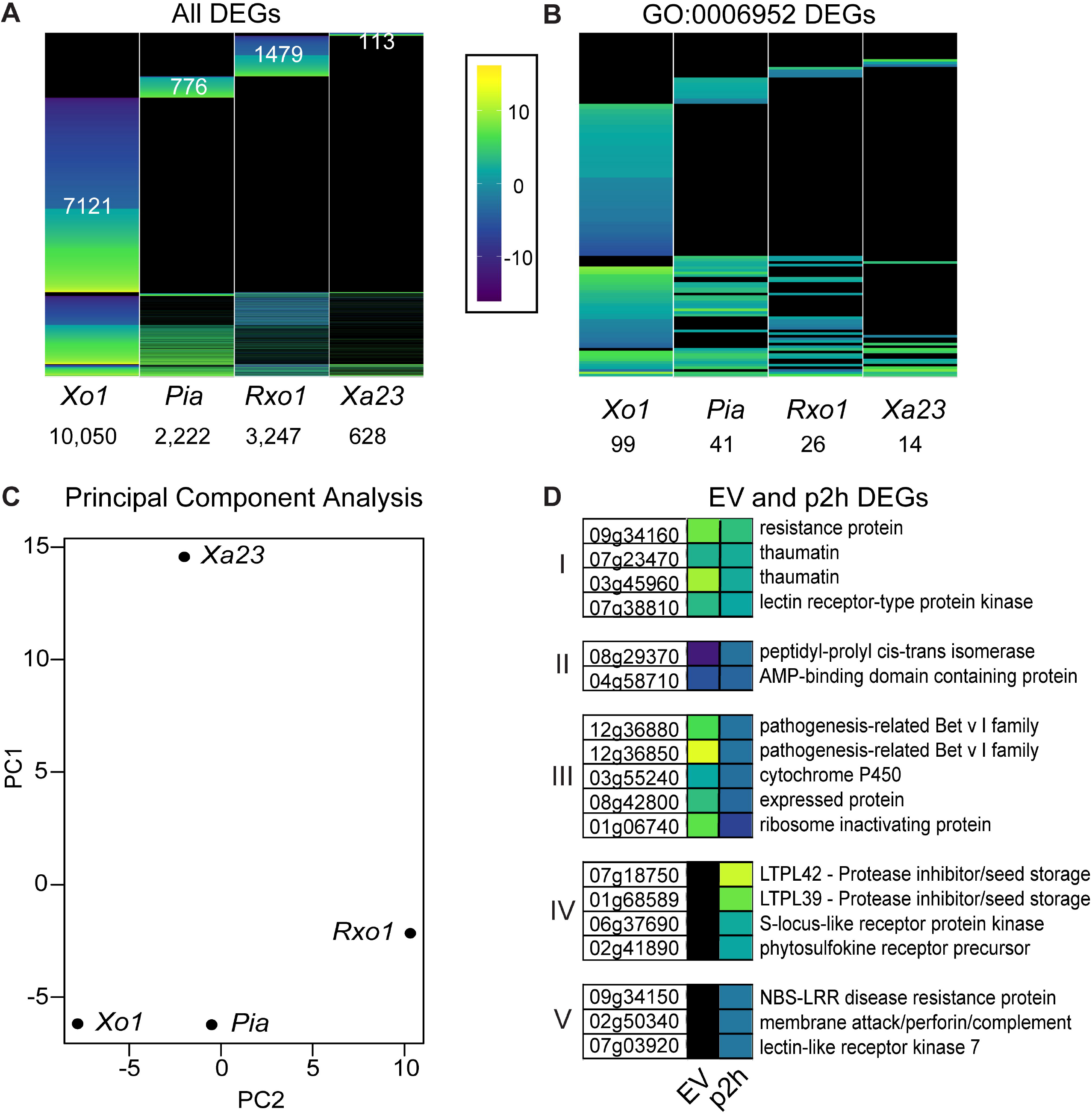
Transgenic Nipponbare plants expressing Xo1_11_ are resistant to African Xoc strain CFBP7331 and the resistance suppressed by a truncTALE. Susceptible cultivar Nipponbare was transformed with pAR902, and leaves of T0 plants from two events were syringe-infiltrated with African Xoc strain CFBP7311 carrying either empty vector (EV) or *tal2h* (p2h) adjusted to OD_600_ 0.4. Leaves were photographed on a light box at 4 days after inoculation. Resistance is apparent as HR (necrosis) at the site of inoculation and disease as expanded, translucent watersoaking.

NLR protein activation is characteristically followed by a suite of responses that includes massive transcriptional reprogramming leading both to HR and to activation of a large number of defense-associated genes (Cui et al., 2015). To gain insight into the nature of *Xo1*-mediated resistance, we compared the global profile of differentially expressed genes during *Xo1*-mediated defense to those of two other NLR genes in rice and to the profile associated with an executor gene. We used our previously reported RNAseq data from Carolina Gold plants inoculated with CFBP7331(EV) or mock inoculum (Read et al., 2020), data for the NLR gene *Pia* for resistance to the rice blast pathogen *Magnaporthe oryzae* (Tanabe et al., 2014), data from rice resistant to bacterial leaf streak due to transgenic expression of the maize NLR gene *Rxo1* (Xie et al., 2007; Zhou et al., 2010), and data for the transcriptomic response associated with induction of the executor resistance gene *Xa23* by an Xoo strain with the corresponding TALE (Tariq et al., 2018). Though limited, these datasets include the only currently available expression data for NLR and executor gene-mediated resistance to *Xanthomonas* in rice. Differentially expressed genes (log_2_-fold change >1 or <−1; *p*-value >0.05) in the comparison between pathogen-inoculated and mock-inoculated plants were compared across the four datasets. The total number of DEGs ranged from 10,050 for *Xo1* to 628 for *Xa23*, and the overall profiles were largely distinct (**Fig. 2A**, **Table S1**). For each resistance gene, there were a number of DEGs found only in the pathogen to mock comparison for that dataset, and this was highest for *Xo1* (7,121 genes) (**Fig. 2A**, **Table S1**). Differences among the overall DEG profiles may be influenced by the expression assay (RNAseq vs. microarray), pathogen, annotation, or timepoints used. To compare the responses further the expression of 340 rice genes associated with plant defense response (gene ontology group 0006952) was examined. The *Xo1* profile comprised the largest number of plant defense DEGs (99) and had more DEGs in common with the other NLR-mediated responses (16 with *Rxo1* and 26 with *Pia*) than with the executor gene response (8) (Fig. 2B). Additionally, each of the NLR-mediated responses resulted in a larger number of differentially expressed defense genes (26 for *Rxo1*, 41 for *Pia*) than the *Xa23* response (14), and based on principle component analysis of the defense DEG profiles, were more similar to one another than to the executor gene response (**Fig. 2B and C** and **Table S2**).

**Fig. 2.**
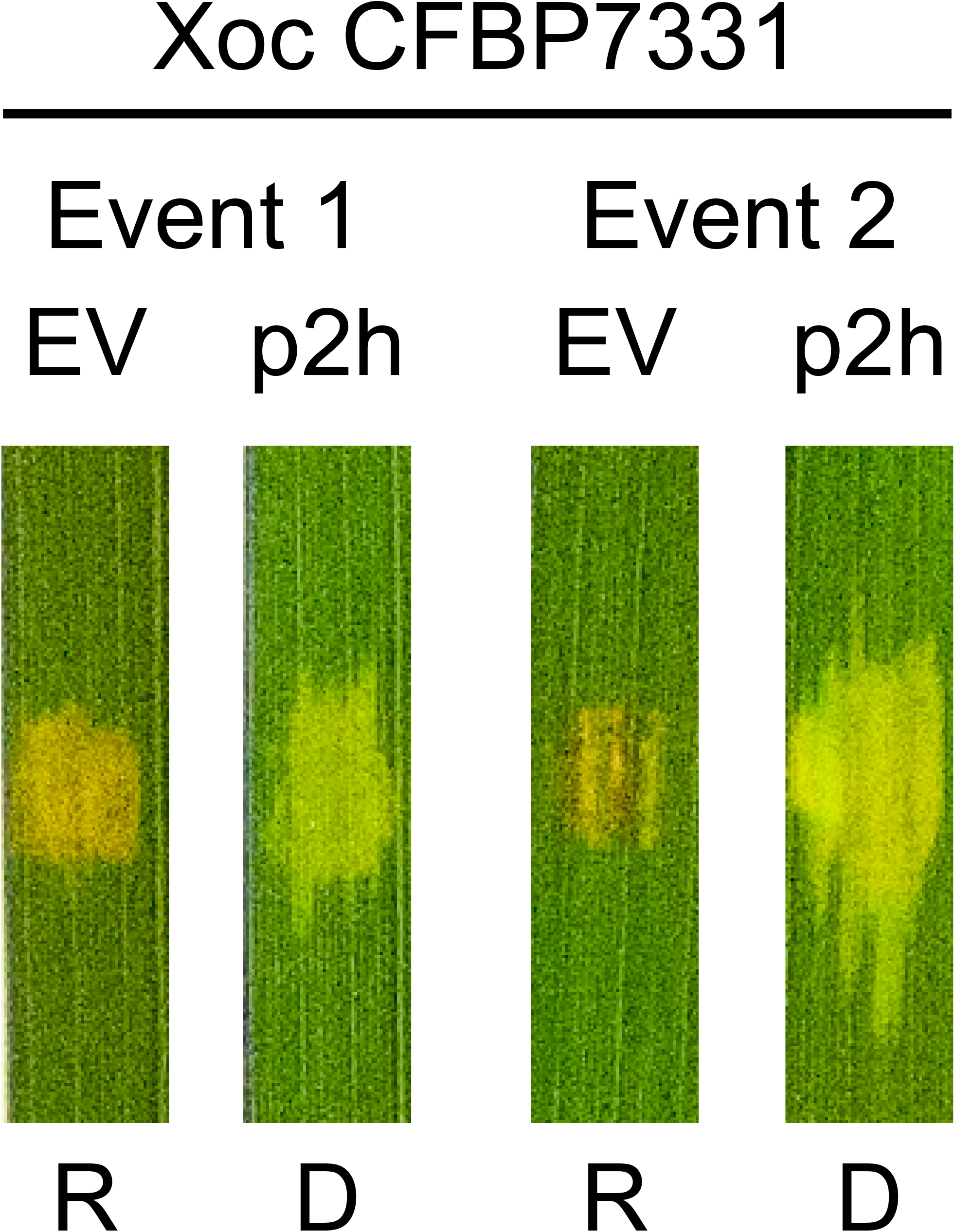
The Xo1-mediated transcriptomic response is similar to those of other NLR genes and is essentially eliminated by Tal2h. **A**, Expression heatmaps (columns) showing all differentially expressed genes (DEGs) in plants undergoing the resistant response compared to mock inoculated plants for *Xo1*, the NLR genes *Pia* and *Rxo1*, and the executor resistance gene *Xa23*. White numbers for each on the heatmap indicate the number of DEGs specific to each response (see **Table S1**). Total numbers of DEGs are indicated below. **B,** Heatmaps for the subset of DEGs from (A) that belong to gene ontology group 0006952, defense response, with totals displayed at bottom. **C**, Principal component analysis. The first two principal components (PC) explain 54.0% and 31.6% of the variation with a total of 85.6%. PC1 demarcated two major clusters: 1) *Xo1*, *Pia*, and *Rxo1*, and 2) *Xa23* **D,** Heatmaps for the 18 defense response DEGs identified in the comparison of Carolina Gold plants inoculated with CFBP7331(p2h) to mock inoculated plants. The “EV” heatmap shows their expression relative to mock in Carolina Gold plants inoculated with CFBP7331(EV) (resistance), and the “p2h” column shows their expression relative to mock in the presence of Tal2h (disease). The DEGs have been divided into five categories: **I**, induced in both; **II**, down-regulated in both; **III**, induced in resistance and down-regulated in disease; **IV**, not differentially expressed in resistance and induced in disease; and **V**, not differentially expressed in resistance and down-regulated in disease.

We also compared DEGs relative to mock in Carolina Gold plants inoculated with CFBP7331(EV) and Carolina Gold plants inoculated with CFBP7331(p2h) (Read et al., 2020), to gain insight into how *Xo1*-mediated resistance is overcome by a pathogen delivering a truncTALE. In contrast to the 99 defense response genes differentially expressed in response to CFBP7331(EV), only 18 defense genes were differentially expressed in response to CFBP7331(p2h) (**Fig. 2C**). Of these 18 genes, 7 were differentially expressed only in the response to the strain with *tal2h*, 4 up and 3 down. Of the remaining 11, 4 were up and 2 were down in both responses, but each less so in the response to the strain with *tal2h*. The other 5 moved in opposite directions entirely, up in the absence but repressed in the presence of *tal2h*, relative to mock. This expression profile during suppression of *Xo1*-mediated resistance is consistent with Tal2h functioning early in the defense cascade. The bacterial leaf streak susceptibility gene *OsSULTR3;6* (Cernadas et al., 2014), activated by Tal8e of CFBP7331 (Wilkins et al., 2015), is strongly induced by both CFBP7331(EV) and CFBP7331(p2h), indicating that TALE function is not compromised by Xo1 or by Tal2h.

The observation that Xo1 reprograms transcription of canonical defense genes upon recognition of the cognate pathogen effector and that reprogramming by Xo1 is essentially blocked by Tal2h led us to explore whether Xo1 localizes to the same subcellular location as TALEs and truncTALEs. Some, but not all, NLR proteins nuclear localize (Shen et al., 2007; Wirthmueller et al., 2007; Caplan et al., 2008; Cheng et al., 2009), and we previously identified putative nuclear localization signals (NLSs) in Xo1_11_ (Read et al., 2020). We generated expression constructs for a green fluorescent protein (GFP) fusion to the N-terminus of Xo1 as well as an N-terminal monomeric red fluorescent protein (mRFP) fusion both to a TALE (Tal1c of Xoc BLS256) and to Tal2h. These constructs were delivered into *Nicotiana benthamiana* leaves using *A. tumefaciens* strain GV3101, and the leaves imaged with a Zeiss 710 confocal microscope (**Fig. 3**). GFP-Xo1 in the absence of either effector but with free mRFP localized to foci that appeared to be nuclei. Co-expression with mRFP-Tal1c or with mRFP-Tal2h confirmed that these foci were nuclei.

**Fig. 3.**
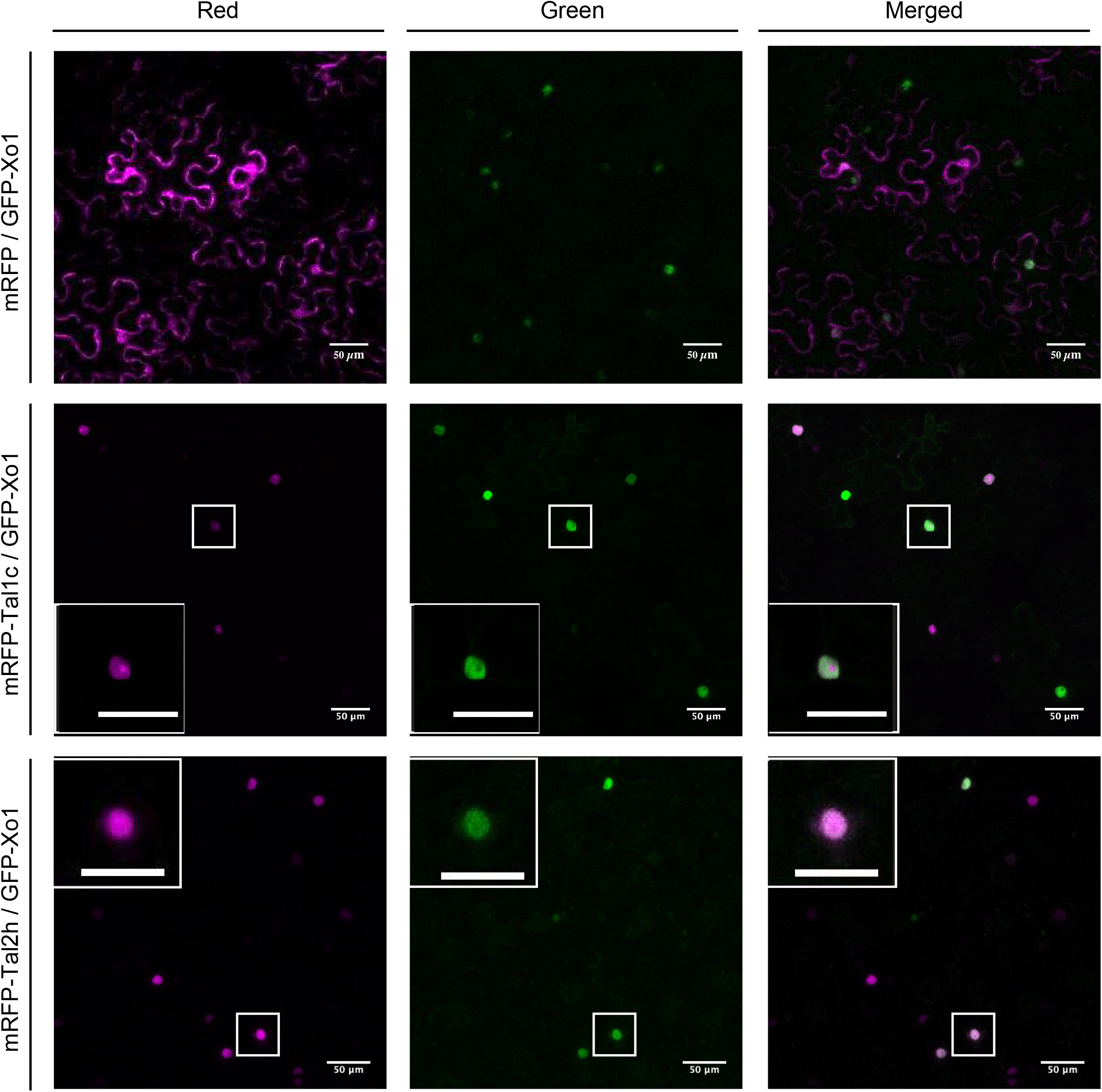
Xo1 localizes to the nucleus. Using *Agrobacterium* co-infiltrations, an expression construct for Xo1 with GFP at the N-terminus (GFP-Xo1) together with a p19 silencing suppressor construct were introduced into *Nicotiana benthamiana* leaves alone or with a construct for mRFP, mRFP fused to TALE Tal1c (mRFP-Tal1c), or mRFP fused to the truncTALE Tal2h (mRFP-Tal2h). Confocal image stacks were taken at 3 days after inoculation and are presented as maximum intensity projections. Insets are magnifications of individual nuclei. The scale bars represent 50 μm.

The localization of Xo1, the TALE, and the truncTALE to the nucleus when transiently expressed in *N. benthamiana* led us to pursue the hypothesis that Xo1 physically interacts with one or both of these proteins in the native context. We generated plasmid constructs that add a 3x FLAG tag to the C-terminus of TALE Tal1c or the truncTALE Tal2h (Tal1c-FLAG and Tal2h-FLAG) and introduced them individually into the TALE-deficient *X. oryzae* strain X11-5A (Triplett et al., 2011) for co-immunoprecipitation from inoculated Carolina Gold leaves (**Fig. 4**). Abilities of the tagged TALE and truncTALE to respectively trigger and suppress *Xo1*-mediated resistance were confirmed (**Fig. S2**). We included also a plasmid for expression of a second, untagged TALE (Tal3c from BLS256) and a plasmid for untagged Tal2h. By pairing the X11-5A transformants with each other or with the untransformed control strain, we were able to probe for Carolina Gold proteins interacting with the tagged TALE or truncTALE, and for interactions of these proteins with each other or with the second TALE. Select combinations were inoculated to Nipponbare leaves for comparison. Inoculation was done by syringe infiltration, in 30-40 contiguous spots on each side of the leaf midrib. For each co-inoculation, tissue was harvested at 48 hours and ground in liquid N_2_, then soluble extract was incubated with anti-FLAG agarose beads and washed to immunopurify the tagged and interacting proteins. Immunoprecipitates were eluted, and an aliquot of each was subjected to western blotting with anti-TALE antibody (**Fig. S3**). The remainders were then resolved on a 4-20% SDS-PAGE and eluates from gel slices containing proteins between approximately 60 and 300 kDa (**Fig. S4**) were digested and the peptides analyzed by mass spectrometry. Proteins were considered present in a sample if at least three peptides mapped uniquely to any of the pertinent annotated genomes searched: the *X. oryzae* strain X11-5A genome (Triplett et al., 2011) plus the TALE(s) or TruncTALE being expressed, the Nipponbare genome (MSU 7; Kawahara et al., 2013), and the Carolina Gold genome (Read et al., 2020). For the Carolina Gold genome, we re-annotated using the RNAseq data from CFBP7331(EV), CFBP7331(p2h), and mock-inoculated plants cited earlier. We carried out the experiment twice.

**Fig. 4.**
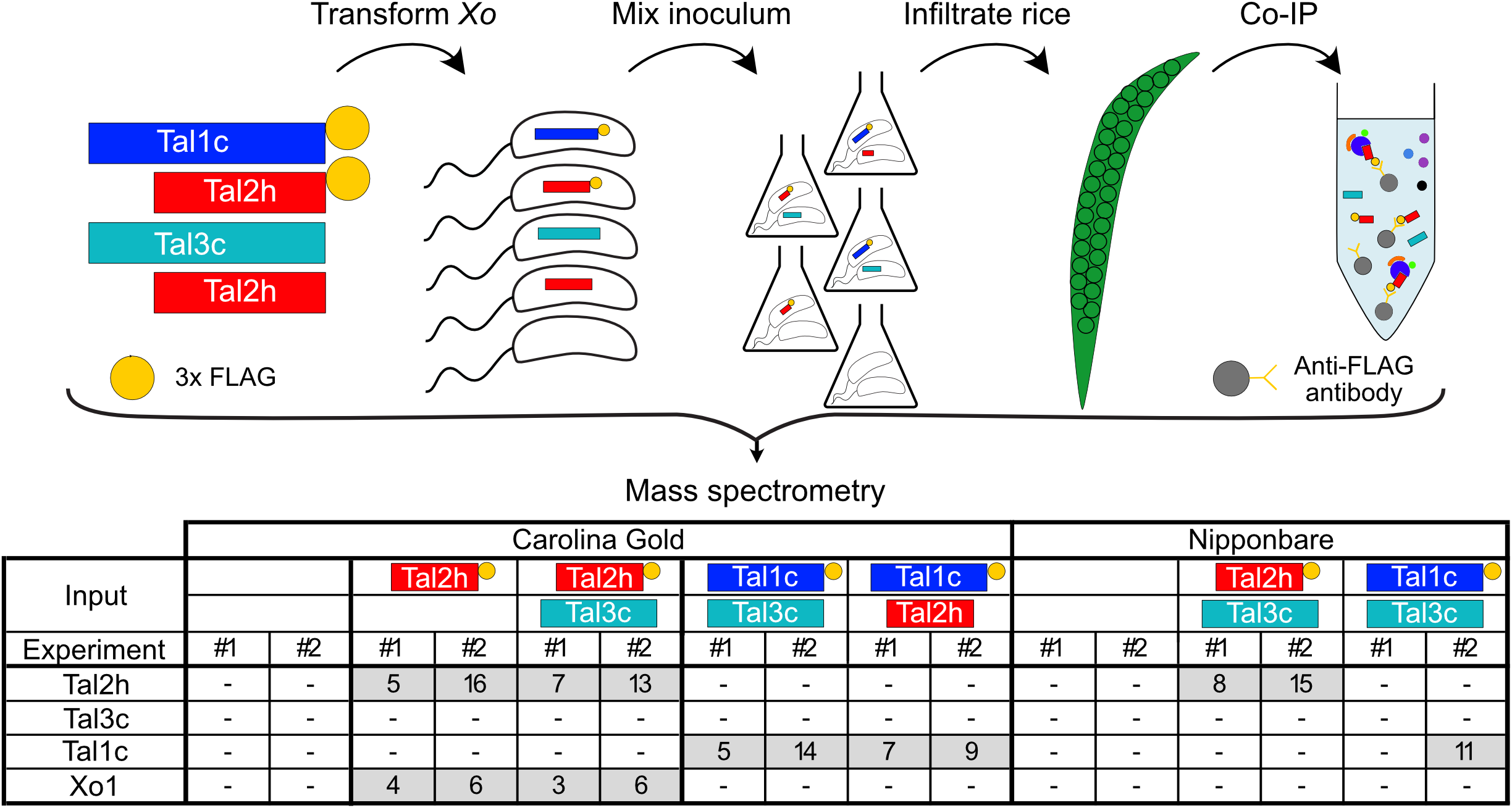
Xo1 co-immunoprecipitates with Tal2h. **Top,** strategy used for co-immunoprecipitation (Co-IP) of truncTALE Tal2h or TALE Tal1c and any interactors. Plasmid borne expression constructs for Tal2h or Tal1c with a C-terminal 3x FLAG tag, as well as untagged Tal2h and a second TALE,Tal3c were introduced into *Xanthomonas oryzae* strain X11-5. Paired combinations of the transformants with each other or with the untransformed control strain, or the control strain alone, were co-infiltrated into leaves of rice varieties Carolina Gold and Nipponbare at a final OD_600_ 0.5 for each transformant. Samples were collected 48 hours after inoculation, ground, and sonicated before Co-IP using anti-FLAG agarose beads. After elution and SDS-PAGE separation, proteins between approximately 60 and 300 kDa were eluted, digested and analyzed by mass spectrometry. The experiment was conducted twice. **Bottom**, co-IP results. For each immunoprecipitate, the numbers of unique peptides detected that matched Tal2h, Tal3c, Tal1c, or Xo1 in each experiment are shown. “-” indicates that ≤ 2 unique peptides were detected.

In the western blot for each experiment (**Fig. S3**), we detected the tagged TALE or truncTALE in each corresponding sample, with the exception of a Tal1c-FLAG/Tal3c/Nipponbare sample in the first experiment. No Tal3c or untagged Tal2h was detected in any sample. The mass spectrometry confirmed these observations, suggesting that neither TALEs with truncTALEs nor TALEs with other TALEs interact appreciably (**Fig. 4**). Xo1 was consistently detected in the Carolina Gold/Tal2h-FLAG samples, irrespective of any co-delivered Tal1c or Tal3c, and not in the Tal1c-FLAG samples or any other sample (**Fig. 4**). No other protein consistently co-purified with Tal2h-FLAG or Tal1c-FLAG in either Carolina Gold or Nipponbare samples (**Dataset S1**).

In summary, we have shown that 1) an NLR protein gene at the *Xo1* locus, harboring an integrated zfBED domain, is *Xo1*; 2) the *Xo1*-mediated response is more similar to those mediated by two other NLR resistance genes than it is to the response associated with TALE-specific transcriptional activation of an executor resistance gene; 3) a truncTALE abolishes or dampens activation of defense-associated genes by *Xo1*; 4) the Xo1 protein, like TALEs and truncTALEs, localizes to the nucleus, and 5) Xo1 specifically co-immunoprecipitates from rice leaves with a pathogen-delivered, epitope-tagged truncTALE. Thus, *Xo1* is an allele or paralog of *Xa1*, and suppression of Xo1 function by a truncTALE is likely the result of physical interaction between the resistance protein and the effector. The latter prediction is consistent with the Xo1 DEG profile during suppression by Tal2h, which suggested that Tal2h functions early in the defense cascade, perhaps by blocking TALE recognition by Xo1.

Whether the interaction between Tal2h and Xo1 is direct or indirect is not certain, but the fact that no other protein was detected consistently that co-immunoprecipitated with Tal2h and Xo1 suggests the interaction is direct. It is tempting to speculate also that TALEs trigger Xo1-mediated resistance by direct interaction with the protein and that truncTALEs function by disrupting the association. Though Tal1c did not pull down Xo1, this might be explained by its lower apparent abundance, based on the western blots. Tal1c might interact weakly or transiently with Xo1, or any complex of the proteins in the plant cells may have begun to degrade with the developing HR at the 48 hour time point sampled. It is also possible that Tal2h interacts with TALEs and masks them from the resistance protein, but both our co-immunoprecipitation results and the fact that Tal2h does not impact TALE activation of the *OsSULTR3;6* susceptibility suggest that this is not the case. An alternative hypothesis is that Xo1 recognition of TALEs is not mediated by a direct interaction between the two proteins.

The results presented constitute an important step toward understanding how Xo1 works, and how its function can be suppressed by the pathogen. Toward determining the relationship of the interaction to defense suppression, an immediate next step might be structure function analysis of the interaction to determine the portion(s) of Xo1 and Tal2h involved. For Xo1, the LRR may be the determinative interacting domain. Our previous comparison of the motifs present in Xo1_11_, Xa1, and the closest Nipponbare homolog (Nb-xo1_5_, which is expressed) revealed that the zfBED and CC domains are identical and the NB-ARC domains nearly so (Read et al., 2020). In contrast, the leucine rich repeat domain of Nb-xo1_5_ differs markedly from those of Xo1 and Xa1, which, with the exception of an additional repeat in Xa1, are very similar. Supporting this hypothesis, differences in the LRR determine the pathogen race specificities of some flax rust resistance genes (Ellis et al., 1999). More broadly, the ability of tagged Tal2h to pull down Xo1 suggests that effector co-immunoprecipitation may be an effective approach to characterizing pathogen recognition mechanisms of other resistance proteins, or for identifying a resistance gene *de novo*.

While this paper was under review, Ji and colleagues (Ji et al., 2020) presented the cloning and functional characterization of several *Xa1* homologs, which also demonstrated that *Xo1_11_* is *Xo1*.

## Acknowledgments

The authors thank M. Carter and B. Szurek for critical reading of the manuscript, Matthew Willmann and the Plant Transformation Facility of Cornell’s School of Integrative Plant Science for carrying out the rice transformation, Sandra Harrington and Susan McCouch for assistance growing the regenerants, and Ruchika Bhawal and Elizabeth Anderson at the Proteomics Facility of the Biotechnology Resource Center at the Cornell University’s Institute of Biotechnology (BRC) for conducting the mass spectrometry. Confocal microscopy was carried out at the BRC’s Imaging Facility. This work was supported by the Plant Genome Research Program of the National Science Foundation (IOS-1444511 to AB), the National Institute of Food and Agriculture of the U.S. Department of Agriculture (2018-67011-28025 to AR), and the Gatsby Charitable Foundation (to MM). We also acknowledge support from the National Institutes of Health to the Proteomics Facility for the Orbitrap Fusion mass spectrometer (shared instrumentation grant 1S10 OD017992-01) and to the Imaging Facility for the Zeiss LSM 710 confocal microscope (shared instrumentation grant S10RR025502).

## Author contributions

AR, MH, FR, and AB conceived and designed the study; AR, MH, and FR carried out the experiments; AR, MH, FR, MM, and AB analyzed data; AR, MH, and AB wrote the manuscript.

## Supplemental files

### 1. Supplemental text and figures

**Materials and methods**

**Fig. S1.** Confirmation of CFBP7331(EV) and CFBP(p2h) inoculum on Nipponbare and Carolina Gold plants.

**Fig. S2.** Symptoms on Carolina Gold and Nipponbare leaves caused by inoculum used for the co-IP experiments.

**Fig. S3.** Western blot of immunoprecipitates using anti-TALE antibody.

**Fig. S4.** SDS-PAGE of immunoprecipitates and size range excised for mass spectrometry.

**Supplemental references**

### 2. Supplemental tables

**Table S1.** DEGs in Fig. 2A (all DEGS)

**Table S2.** DEGs in Fig. 2B (GO:0006952 DEGs)

**Table S3.** DEGs in Fig. 2C (GO:0006952 in disease)

### 3. Dataset S1

Mass spectrometry data

## Notes

### Competing Interest Statement

The authors have declared no competing interest.

### Summary of Updates

Some improvements following an initial review. An error in figure 2 and figure 2 legend was corrected. Sections were re-written to make clear the lack of evidence for physical characterization between TALEs and Xo1. Figure 4 was modified for clarity. A principal component analysis was added to quantify relationships among expression datasets (Fig 2).

## References

Bogdanove, A.J., Schornack, S., and Lahaye, T. 2010. TAL effectors: finding plant genes for disease and defense. Curr. Opin. Plant Biol. 13:394–401.

Caplan, J.L., Mamillapalli, P., Burch-Smith, T.M., Czymmek, K., and Dinesh-Kumar, S.P. 2008. Chloroplastic protein NRIP1 mediates innate immune receptor recognition of a viral effector. Cell 132:449–462.

Cernadas, R.A., Doyle, E.L., Nino-Liu, D.O., Wilkins, K.E., Bancroft, T., Wang, L., Schmidt, C.L., Caldo, R., Yang, B., White, F.F., Nettleton, D., Wise, R.P., and Bogdanove, A.J. 2014. Code-assisted discovery of TAL effector targets in bacterial leaf streak of rice reveals contrast with bacterial blight and a novel susceptibility gene. PLoS Path. 10:e1003972.

Cheng, Y.T., Germain, H., Wiermer, M., Bi, D., Xu, F., Garcia, A.V., Wirthmueller, L., Despres, C., Parker, J.E., Zhang, Y., and Li, X. 2009. Nuclear pore complex component MOS7/Nup88 is required for innate immunity and nuclear accumulation of defense regulators in Arabidopsis. Plant Cell 21:2503–2516.

Cui, H., Tsuda, K., and Parker, J.E. 2015. Effector-triggered immunity: from pathogen perception to robust defense. Annu. Rev. Plant Biol. 66:487–511.

Ellis, J.G., Lawrence, G.J., Luck, J.E., and Dodds, P.N. 1999. Identification of regions in alleles of the flax rust resistance gene *L* that determine differences in gene-for-gene specificity. Plant Cell 11:495–506.

Hutin, M., Perez-Quintero, A.L., Lopez, C., and Szurek, B. 2015. MorTAL Kombat: the story of defense against TAL effectors through loss-of-susceptibility. Front. Plant Sci. 6:535.

Ji, C., Ji, Z., Liu, B., Cheng, H., Liu, H., Liu, S., Yang, B., and Chen, G. 2020. *Xa1* allelic *R* genes activate rice blight resistance suppressed by Interfering TAL effectors. Plant Communications:100087.

Ji, Z., Ji, C., Liu, B., Zou, L., Chen, G., and Yang, B. 2016. Interfering TAL effectors of *Xanthomonas oryzae* neutralize *R*-gene-mediated plant disease resistance. Nat. Commun. 7:13435.

Kawahara, Y., de la Bastide, M., Hamilton, J.P., Kanamori, H., McCombie, W.R., Ouyang, S., Schwartz, D.C., Tanaka, T., Wu, J., and Zhou, S. 2013. Improvement of the *Oryza sativa* Nipponbare reference genome using next generation sequence and optical map data. Rice 6:4.

Lolle, S., Stevens, D., and Coaker, G. 2020. Plant NLR-triggered immunity: from receptor activation to downstream signaling. Curr. Opin. Immunol. 62:99–105.

McClung, A., and Fjellstrom, R. 2010. Using molecular genetics as a tool to identify and refine “Carolina Gold”. Pages 37–41 in: The Golden Seed: Writings on the History and Culture of Carolina Gold Rice, D.S. Shields, ed. Douglas W. Bostick for the Carolina Gold Rice Foundation.

Perez-Quintero, A.L., and Szurek, B. 2019. A decade decoded: spies and hackers in the history of TAL effectors research. Annu. Rev. Phytopathol. 57:459–481.

Read, A.C., Rinaldi, F.C., Hutin, M., He, Y.Q., Triplett, L.R., and Bogdanove, A.J. 2016. Suppression of *Xo1*-mediated disease resistance in rice by a truncated, non-DNA-binding TAL effector of *Xanthomonas oryzae*. Front. Plant Sci. 7:1516.

Read, A.C., Moscou, M.J., Zimin, A.V., Pertea, G., Meyer, R.S., Purugganan, M.D., Leach, J.E., Triplett, L.R., Salzberg, S.L., and Bogdanove, A.J. 2020. Genome assembly and characterization of a complex zfBED-NLR gene-containing disease resistance locus in Carolina Gold Select rice with Nanopore sequencing. PLoS Genet. 16:e1008571.

Shen, Q.H., Saijo, Y., Mauch, S., Biskup, C., Bieri, S., Keller, B., Seki, H., Ulker, B., Somssich, I.E., and Schulze-Lefert, P. 2007. Nuclear activity of MLA immune receptors links isolate-specific and basal disease-resistance responses. Science 315:1098–1103.

Tanabe, S., Yokotani, N., Nagata, T., Fujisawa, Y., Jiang, C., Abe, K., Ichikawa, H., Mitsuda, N., Ohme-Takagi, M., Nishizawa, Y., and Minami, E. 2014. Spatial regulation of defense-related genes revealed by expression analysis using dissected tissues of rice leaves inoculated with *Magnaporthe oryzae*. J. Plant Physiol. Pathol. 2:1000135.

Tariq, R., Wang, C., Qin, T., Xu, F., Tang, Y., Gao, Y., Ji, Z., and Zhao, K. 2018. Comparative transcriptome profiling of rice near-isogenic line carrying *Xa23* under infection of *Xanthomonas oryzae* pv. oryzae. Int. J. Mol. Sci. 19:717.

Triplett, L.R., Hamilton, J.P., Buell, C.R., Tisserat, N.A., Verdier, V., Zink, F., and Leach, J.E. 2011. Genomic analysis of *Xanthomonas oryzae* isolates from rice grown in the United States reveals substantial divergence from known *X. oryzae* pathovars. Appl. Environ. Microbiol. 77:3930–3937.

Triplett, L.R., Cohen, S.P., Heffelfinger, C., Schmidt, C.L., Huerta, A.I., Tekete, C., Verdier, V., Bogdanove, A.J., and Leach, J.E. 2016. A resistance locus in the American heirloom rice variety Carolina Gold Select is triggered by TAL effectors with diverse predicted targets and is effective against African strains of *Xanthomonas oryzae* pv. oryzicola. Plant J. 87:472–483.

Wilkins, K.E., Booher, N.J., Wang, L., and Bogdanove, A.J. 2015. TAL effectors and activation of predicted host targets distinguish Asian from African strains of the rice pathogen *Xanthomonas oryzae* pv. oryzicola while strict conservation suggests universal importance of five TAL effectors. Front. Plant Sci. 6:536.

Wirthmueller, L., Zhang, Y., Jones, J.D., and Parker, J.E. 2007. Nuclear accumulation of the Arabidopsis immune receptor RPS4 is necessary for triggering EDS1-dependent defense. Curr. Biol. 17:2023–2029.

Xie, X.W., Yu, J., Xu, J.L., Zhou, Y.L., and Li, Z.K. 2007. [Introduction of a non-host gene *Rxo1* cloned from maize resistant to rice bacterial leaf streak into rice varieties]. Sheng Wu Gong Cheng Xue Bao 23:607–611.

Yoshimura, S., Yamanouchi, U., Katayose, Y., Toki, S., Wang, Z.X., Kono, I., Kurata, N., Yano, M., Iwata, N., and Sasaki, T. 1998. Expression of *Xa1*, a bacterial blight-resistance gene in rice, is induced by bacterial inoculation. Proc. Natl. Acad. Sci. USA 95:1663–1668.

Zhang, J., Yin, Z., and White, F. 2015. TAL effectors and the executor *R* genes. Front. Plant Sci. 6:641.

Zhou, Y.L., Xu, M.R., Zhao, M.F., Xie, X.W., Zhu, L.H., Fu, B.Y., and Li, Z.K. 2010. Genome-wide gene responses in a transgenic rice line carrying the maize resistance gene *Rxo1* to the rice bacterial streak pathogen, *Xanthomonas oryzae* pv. oryzicola. BMC Genomics 11:78.

