## Supplemental text and figures for "Cloning of the rice *Xo1* resistance gene and interaction of the Xo1 protein with the defense-suppressing *Xanthomonas* effector Tal2h"

**Materials and methods**

**Fig. S1.** Confirmation of CFBP7331(EV) and CFBP(p2h) inoculum on Nipponbare and Carolina Gold plants.

**Fig. S2.** Symptoms on Carolina Gold and Nipponbare leaves caused by inoculum used for the co-IP experiments.

**Fig. S3.** Western blot of immunoprecipitates using anti-TALE antibody.

**Fig. S4.** SDS-PAGE of immunoprecipitates and size range excised for mass spectrometry.

**Supplemental references**

### Supplemental Materials and methods

#### *Generation and testing of $Xo1_{11}$ transgenic Nipponbare*

The binary transformation plasmid pAR902, containing the *$Xo1_{11}$* promoter sequence, a Kozak sequence, the genomic *$Xo1_{11}$* gene body, and a 35S terminator, was generated through several steps. Genomic *$Xo1_{11}$* was amplified in two parts using oligo pairs 2492/2420 and 2421/2489. Amplicons were ligated into backbone vector pDONR221 following digestion with *Not*I, *Sph*I, and *Asc*I. This cloning strategy resulted in an undesired frameshift downstream of the coding sequence that made the clone incompatible with Gateway destination vectors. The frameshift was corrected by amplifying a small fragment with oligo pair 2644/2645 then ligating it into the *Cla*I/*Asc*I digested parent plasmid. A Kozak consensus sequence was introduced by QuickChange (Agilent) using primer pair 3267/3268. The 993 bp region upstream of the *$Xo1_{11}$* start codon was amplified with oligo pair 3080/3082 and BP Gateway cloned into pDONR P4P1r (ThermoFisher). The plasmids containing the promoter sequence, the genomic *$Xo1_{11}$* sequence, and a pDONR P2rP3 with a 35S terminator were LR Gateway cloned into the binary Gateway destination plasmid pKm43GW (Karimi et al., 2005).

Sequence confirmed pAR902 was electroporated into *Agrobacterium tumefaciens* strain EHA101 for transformation of japonica rice cultivar Nipponbare, which was performed by the Cornell University Plant Transformation Facility.

T0 regenerants were transferred to 4 inch pots containing LC-1 soil mixture (Sungro) and moved to a PGC15 (Percival Scientific) growth chamber ~60 cm below a combination of fluorescent and incandescent bulbs providing ~1000 µmoles/$m^2$/s measured at 15 cm, under a cycle of 12 h light at 28°C and 12 h dark at 25°C. Wildtype Nipponbare and Carolina Gold Select seeds were sown to serve as bacterial inoculation controls.

*Xanthomonas oryzae* pv. oryzicola strain CFBP7331 transformed with empty vector (pAC99; Cernadas et al., 2014) or p2h (Read et al., 2016) were grown overnight in liquid GYE with appropriate antibiotics at 28°C with shaking. Overnight cultures were pelleted, washed once, and resuspended in 10 mM $MgCl_2$ to an $OD_{600}$ of 0.4 prior to infiltration using a needleless syringe (Reimers et al., 1992). Inoculated plants were maintained in the growth chamber under the above described conditions. Leaves were photographed on a light box.

#### *Gene expression analysis*

For *Xo1*, RNAseq data for mock-inoculated (SRR9320033, SRR9320038, SRR9320039), CFBP7331(EV)-inoculated (SRR9320035, SRR9320036, SRR9320037) and CFBP7331(p2h)-inocualted (SRR9320034, SRR9320040, SRR9320041) Carolina Gold Select plants (Read et al., 2020) were downloaded from the NCBI Sequence Read Archive. Reads were aligned to the MSU v7 Nipponbare reference genome annotation using the "—quantMode GeneCounts" option in STAR (Dobin et al., 2013). STAR output was used in DESEQ2 (Love et al., 2014) to determine differentially expressed genes (DEGs) in CFBP7331(EV) and CFBP7331(EV) compared to mock.

For *Pia*, NCBI GEO2R (Smyth, 2004; Davis and Meltzer, 2007) was used to identify DEGs from microarray data of rice line NP/Pia expressing the rice blast resistance gene *Pia* (GEO GSE62893). Those data had been collected 24 hpi following mock-inoculation (GSM1535715, GSM1535731, GSM1535750) or inoculation with avirulent *Magnoporthe oryzae* strain P91-15B, race 001.0 (GSM1535711, GSM1535727, GSM1535743) (Tanabe et al., 2014).

For *Rxo1*, NCBI GEO2R (Smyth, 2004; Davis and Meltzer, 2007) was used to identify DEGs from microarray data of transgenic rice expressing the maize NLR gene *Rxo1* (GEO GSE19239) (Zhou et al., 2010). Data had been collected 48 hpi following mock-inoculation (GSM476772, GSM476773, GSM476774) or inoculation with avirulent Xoc strain FJR5 (GSM476769, GSM476770, GSM476771).

For *Xa23*, DEGs were as reported in a previous RNAseq analysis of rice variety CBB23 inoculated with strain PXO99A compared with mock inoculated CBB23 at 24 hpi (Tariq et al., 2018).

DEGs with $log_2$-fold change between -1 and 1 or with *p*-value >0.05 were excluded from the analysis. Heatmaps were generated in R version 3.6.3 using the package Superheat (Barter and Yu, 2018).

Principal component analysis was performed in R using the log fold-change values of differentially expressed defense genes between inoculated and mock-inoculated samples in the four data sets. In total, 181 genes were used in the analysis.

***Localization of Xo1***

The expression construct for green fluorescent protein fused to the N-terminus of Xo1 was generated by LR Gateway recombination between the $Xo1_{11}$ entry vector and pGWB6 (Nakagawa et al., 2007). Monomeric red fluorescent protein fusions to the N-termini of Tal1c and Tal2h were generated by LR Gateway cloning of *tal1c* and *tal2h* entry plasmids with pGWB455 (Tanaka et al., 2011). Empty pGWB455 was used as the free mRFP control construct. Expression constructs were electroporated into *A. tumefaciens* strain GV3101. Overnight cultures grown at 28°C in LB broth with antibiotic were pelleted and washed in infiltration buffer (10 mM MES, pH 5.6, 10 mM $MgCl_2$, and 150 μM acetosyringone). Washed pellets were resuspended in infiltration buffer, adjusted to $OD_{600}$ 1.0, and mixed such that each strain was present at $OD_{600}$ 0.4. Each mixture also contained GV3101 carrying an expression construct for the silencing suppressor p19 at $OD_{600}$ 0.2. Mixtures were incubated for 4 hours at room temperature before blunt syringe infiltration of 4-week-old *Nicotiana benthamiana* leaves. Plants were left on the bench under fluorescent light for three days. Then, leaf discs were punched from the infiltrated area and observed with a Zeiss 710 confocal microscope at the Cornell Biotechnology Core Facility. GFP was imaged with excitation 488 nm and detection at 498-532 nm, while mRFP was imaged with excitation 561 nm and detection at 606 to 621 nm. Maximum intensity projections were generated in FIJI (Schindelin et al., 2012).

***Co-immunoprecipitation, western blot analysis, and mass spectrometry***

Plasmid pFR339 encoding Tal2h:3xFLAG was constructed by inserting a gBlock fragment at the restriction sites *Aat*II and *EcoR*V in-frame with *tal2h* in the plasmid pAR009_2h (Read et al., 2016) The gBlock encodes a fragment of the C-terminal domain of Tal2h followed by three tandem FLAG epitope tags. Plasmid pFR340 encoding Tal1c:3xFLAG was constructed by inserting a gBlock fragment at the restriction sites *Bbv*CI and *Eco*RV in-frame with *tal1c* in the plasmid pAR009_1c (Read et al., 2016). The gBlock encodes a fragment of the C-terminal domain of Tal1C followed by three tandem FLAG epitope tags. gBlocks were synthesized by Integrated DNA Technologies. pFR339 and pFR340 were used as entry vectors to transfer the coding sequences into *Xanthomonas* expression vector pKEB31 (Cermak et al., 2011) by Gateway LR reaction (ThermoFisher Scientific). These constructs were transformed by electroporation into the TALE-deficient *X.oryzae* strain X11-5A (Triplett et al., 2011).

Nipponbare and Carolina Gold Select plants were grown in a growth chamber under cycles of 12 hours of light at 28°C and 75-80% relative humidity (RH), and 12 hours of dark at 25°C, and inoculated by syringe infiltration (Reimers et al., 1992) at 5 weeks old. For inoculation, individual bacterial suspensions containing each strain were made in 10 mM $MgCl_2$ at an $OD_{600}$ of 1, and mixed at equal volume to obtain the final suspensions for co-infiltration. A total of ten leaves with 60 to 80 tandem infiltration spots per leaf were inoculated for each co-inoculum. Leaves were collected 48 hpi and immediately frozen in liquid nitrogen.

Leaves frozen in liquid nitrogen were finely ground with a mortar and pestle. Three grams of leaf tissue were resuspended in 3 mL of GTEN extraction buffer (10% glycerol, 25 mM Tris pH 7.5, 1 mM EDTA, 150 mM NaCl) with 2% w/v polyvinylpolypyrrolidone, 1x protease inhibitor cocktail (Sigma Aldrich), and 0.1% Tween 20. Insoluble debris was pelleted by centrifugation at 3,000 x g for 20 min at 4°C. Samples were sonicated on ice and a second centrifugation at 20,000 x g for 20min at 4°C was done before transferring the supernatant to a new tube and adjusting to 2 mL with immunoprecipitation buffer (GTEN, with 0,1% Tween 20). EZview Red ANTI-FLAG M2 Affinity gel (Sigma Aldrich) was washed 2 times with 5 volumes of IP buffer and resuspended in the original volume. 5 µl of the suspension was added to each sample and the samples incubated with mixing by turning end-over-end for 2 h at 4°C. Resin was washed seven times in IP buffer, then bound proteins were eluted with 40 µl of 1x SDS Laemmli sample buffer without reducing agent. 1x dithiothreitol (DTT) was then added to each eluate and the samples incubated for 10 min at 95°C.

For western blotting, a 5 µl aliquot of each sample was resolved by 7.5% Tris-Glycine SDS-PAGE (Bio-Rad). Proteins were transferred to a membrane using the Trans-Blot Turbo Transfer System (Bio-Rad). Immunoblotting was performed using standard procedures. TALEs were detected with a 1:5000 diluted primary anti-TALE rabbit polyclonal antibody and a 1:1000 diluted secondary HRP conjugated goat anti-rabbit IgG antibody (Thermo Fisher). TALEs were visualized by chemiluminescence using the Clarity ECL substrate (Bio-Rad). The anti-TALE antibody was raised by Pocono Rabbit Farm and Laboratory (Canadensis, PA) using a 3 mg/ml sample of a dTALE expressed in *E. coli* and affinity-purified as described previously (Rinaldi et al., 2017). The antiserum was tested for specificity, and a dilution of 1:5000 was determined to be optimal for westernblot immunodetection of TALEs. Aliquots were preserved in -80°C in the presence of sodium azide.

Next, 30 µl of eluate from each immunoprecipitate were loaded in a 4-20% Tris Glycine SDS-PAGE stain free gel (Bio-Rad). The gel was stained using SYPRO Ruby protein gel stain (Invitrogen) following manufacturer instructions. The portion of each lane spanning molecular weight markers from 60 kDa to approximately 300 kDa was excised for mass spectrometry analysis. The excised gel fragments were subjected to in-gel digestion followed by extraction of the tryptic peptide as reported previously (Yang et al., 2007). The gel pieces were washed consecutively with 600 µL distilled/deionized water followed by 50 mM ammonium bicarbonate, 50% acetonitrile (ACN), and finally 100% ACN. The dehydrated gel pieces were reduced with 250 µL of 10 mM DTT in 100 mM ammonium bicarbonate for 1 hour at 56 °C, then alkylated with 250 µL of 55 mM iodoacetamide in 100 mM ammonium bicarbonate at room temperature in the dark for 45 minutes.Wash steps were repeated as described above. The gel slices were then dried and rehydrated with trypsin (Promega) at an estimated 1:3 w/w ratio in 50 mM ammonium bicarbonate and 10% ACN and incubated at 37 °C for 18 h. The digested peptides were extracted twice with 200 µl of 50% ACN and 5% formic acid (FA) and once with 200 µl of 75% ACN and 5% FA. For each sample, extracts were combined and filtered with a Costar Spin-X 0.22 µm spin filter (Corning) and dried in a speed vacuum. Each sample was reconstituted in 2% ACN and 0.5% FA prior to LC MS/MS analysis.

Nano LC-ESI-MS/MS analysis was carried out using an Orbitrap Fusion Tribrid mass spectrometer (Thermo-Fisher Scientific) equipped with a nanospray Flex Ion Source and coupled with a Dionex UltiMate 3000 RSLCnano system (Thermo-Fisher Scientific) (Thomas et al., 2017; Yang et al., 2018). For each reconstituted sample, 8 µl were injected onto a PepMap C-18 RP nano trapping column (5 µm, 100 µm i.d x 20 mm) at 15 µL/min flow rate for rapid sample loading and then separated on a PepMap C-18 RP nano column (2 µm, 75 µm x 25 cm) at 35 °C. The tryptic peptides were eluted in a 60 min gradient of 5% to 38% ACN in 0.1% FA at 300 nL/min, followed by a 7 min ramping to 90% ACN-0.1% FA and an 8 min hold at 90% ACN-0.1% FA. The column was re-equilibrated with 0.1% FA for 25 min prior to the next run. The Orbitrap Fusion was operated in positive ion mode with spray voltage set at 1.6 kV and source temperature at 275°C. External calibration for FT, IT and quadrupole mass analyzers was performed. In data-dependent acquisition (DDA) analysis, the instrument was operated using FT mass analyzer in MS scan to select precursor ions followed by 3 sec “Top Speed” data-dependent CID ion trap MS/MS scans at 1.6 m/z quadrupole isolation for precursor peptides with multiple charged ions above a threshold ion count of 10,000 and normalized collision energy of 30%. MS survey scans at a resolving power of 120,000 (fwhm at m/z 200), for the mass range of m/z 375-1575. Dynamic exclusion parameters were set at 40 sec of exclusion duration with ±10 ppm exclusion mass width. All data were acquired under Xcalibur 3.0 operation software (Thermo-Fisher Scientific).

The raw DDA files for CID MS/MS were queried against peptide databases using Proteome Discoverer (PD) 2.2 software (Thermo Fisher Scientific) using the Sequest HT algorithm. The PD 2.2 processing workflow containing an additional node of Minora Feature Detector for precursor ion-based quantification was used for protein identification. The databases were generated from the *X. oryzae* strain X11-5A genome, encoding 3,546 proteins (Triplett et al., 2011), plus Tal1c, Tal2h, and Tal3c; the Nipponbare genome (MSU 7; Kawahara et al., 2013), containing 47,418 protein entries; and the Carolina Gold Select genome (Read et al., 2020). For the Carolina Gold Select genome, we reannotated by using hisat2 (Kim et al., 2019) to map to the genome the RNAseq reads we generated previously (Read et al., 2020), then cufflinks (Trapnell et al., 2012) to build gene models. This resulted in 33,956 protein entries. We added the

*Xo1*$_{11}$ gene model we predicted previously based on comparative structural analysis (Read et al., 2020). To generate the peptide database, two-missed trypsin cleavage sites were allowed. The peptide precursor tolerance was set to 10 ppm, and fragment ion tolerance was set to 0.6 Da. For the database search, variable modification of methionine oxidation, deamidation of asparagines/glutamine, and fixed modification of cysteine carbamidomethylation were set. Only high confidence peptides defined by Sequest HT with a 1% FDR by Percolator were considered for the peptide identification. The final protein IDs contained protein groups that were filtered with at least 2 peptides per protein. The precursor abundance intensity for each peptide identified by MS/MS in each sample was automatically determined and the unique peptides for each protein in each sample were summed and used for calculating the protein abundance by PD 2.2 software without normalization.

***Plasmids used***

**Cloning *Xo1***

| Plasmid | Description | Purpose | Reference |
|---|---|---|---|
| pKM43GW | Binary Gateway destination vector, $Km^r$ | destination | (Karimi et al., 2005) |
| pAR902 | Binary Gateway destination vector pKM43GW containing the *Xo1*$_{11}$ promoter, *Xo1*$_{11}$ sequence of pDONR221:KozXo1, and 35S terminator, $Sp^r$ | transformation | This study |
| pDONR221:KozXo1 | pDONR221 entry vector containing a 'CACC' Kozak sequence immediately upstream of the genomic sequence of *Xo1*$_{11}$ with stop codon deleted, $Km^r$ | intermediate | This study |
| Intermediate:Xo1_1 | Genomic clone of *Xo1*$_{11}$ with stop codon deleted. Note that cloning strategy resulted in frame-shift for any C-terminal fusions and construct does not include Kozak sequence, $Km^r$ | intermediate | This study |
| Intermediate:Xo1_2 | Same as Intermediate:Xo1_1, but C-terminal frame-shift was corrected, $Km^r$ | intermediate | This study |
| pDONR41:Xo1Promoter | pDONR_P4-P1r containing the 1kb genomic sequence upstream of *Xo1*$_{11}$, $Km^r$ | intermediate | This study |
| pDONR_P4-P1r | donor vector with attP4 and attP1 (reversed) sites, $Km^r$, $Cm^r$ | intermediate | Invitrogen |
| pDONR23:35St | pDONR_P2r-P3 containing an in-frame stop codon followed by a 35S terminator, $Km^r$ | intermediate | (Ivanov and Harrison, 2014) |
| pDONR_P2r-P3 | donor vector with attP2 (reversed) and attP3 sites, $Km^r$, $Cm^r$ | intermediate | Invitrogen |

**Inoculation assay**

| Plasmid | Description | Purpose | Reference |
|---|---|---|---|
| pKEB31 | pDD62 derivative containing Gateway destination vector cassette (Invitrogen) between *Xba*I and *Bam*HI sites, $Tc^r$ | expression | (Cermak et al., 2011) |
| pAC99 | pKEB31 containing *tal1c* of BLS256 missing the *Sph*I repeat-encoding fragment, $Tc^r$ | expression | (Cernadas et al., 2014) |
| pKEB31-8-2h | pKEB31 containing Gateway fragment of pAR008-2h, $Tc^r$ | expression | (Read et al., 2016) |

**Localization**

| Plasmid | Description | Purpose | Reference |
|---|---|---|---|
| pGWB455 | Binary Gateway destination vector for transformation or transient expression of mRFP-tagged proteins, $Sp^r$, $Cm^r$ | intermediate/ expression | (Nakagawa et al., 2007; Tanaka et al., 2011) |
| pGWB6 | Binary Gateway destination vector for transformation or transient expression of GFP-tagged proteins, $Km^r$, $Cm^r$ | intermediate/ expression | (Nakagawa et al., 2007) |
| pGWB455:Tal1c | Binary vector pGWB455 containing mRFP fusion at N-terminus of Tal1c from pAR009-1c, $Sp^r$ | expression | This study |
| pAR008-2h | pAR008 containing the *Aat*II-*Sph*I CRR of tal2h, $Ap^r$ | intermediate | (Read et al., 2016) |
| pAR009-1c | pAR009 containing the *Aat*II-*Sph*I CRR of tal1c, $Ap^r$ | intermediate | (Read et al., 2016) |
| pGWB455:Tal2h | Binary vector pGWB455 containing mRFP fusion at N-terminus of Tal2h from pAR008-2h, $Sp^r$ | expression | This study |
| pGWB6:Xo1 | Binary vector pGWB6 containing GFP fusion at N-terminus of genomic *Xo1*$_{11}$ sequence from Intermediate:Xo1_2, $Km^r$ | expression | This study |

**Co-IP**

| | | | |
|---|---|---|---|
| pTal1c-3xFLAG | Tal1c sequence from pAR009-1c with C-terminal 3xFLAG tag in pKEB31 expression backbone, $Tc^r$ | expression | This study |
| pTal2h-3xFLAG | Tal2h sequence from pAR008-2h with C-terminal 3xFLAG tag in pKEB31 expression backbone, $Tc^r$ | expression | This study |

***Oligonucleotides used***

| Oligo | Sequence | Purpose |
|---|---|---|
| b2492 | CAAAAAAGCAGGCTCCGCGGCCGCCACCAA AATGGAGGATGTGGAAGCCGGTTTGC | Forward primer for amplifying *Xo1*$_{11}$ genomic sequence - contains NotI site |
| b2420 | TCATTACCAAAAGCATGCACTTTAAATAGTGA | Reverse primer for amplifying *Xo1*$_{11}$ genomic sequence - contains SphI site |
| b2421 | TGGTCACTATTTAAAGTGCATGCTTTTGGTAA | Forward primer for amplifying *Xo1*$_{11}$ genomic sequence - contains SphI site |
| b2489 | AGAAAGCTGGGTCGGCGCGCCGACAATGCA TTGGAGCGGATT | Reverse primer for amplifying *Xo1*$_{11}$ genomic sequence - contains AscI site |
| b2644 | CGACATCGATGACCCCTCTATCC | Forward primer to fix *Xo1*$_{11}$ C-terminus frame-shift - contains ClaI site |
| b2645 | GTCGGCGCGCCCGTTCACATATCCCCCATT AATTTTG | Reverse primer to fix *Xo1*$_{11}$ C-terminus frame-shift - contains AscI site |
| b3267 | CACCATGGAGGAGGTGGAAGCC | Forward primer to add 'CACC' Kozak sequence |
| b3268 | GAAGCCTGCTTTTTTGTACAAAG | Reverse primer to add 'CACC' Kozak sequence |
| b3080 | GGGGACAACTTTGTATAGAAAAGTTGCCTCG AGGGTATAGACCATATTTCCCCTG | Forward primer to amplify and clone *Xo1*$_{11}$ promoter sequence |
| b3082 | GGGGACTGCTTTTTTGTACAAACTTGCGGAA CAGGAGCAGTCCTTGGACTG | Reverse primer to amplify and clone *Xo1*$_{11}$ promoter sequence |
| gBlock 2h | CTGCTTCGGCGGACGTCCTGCCCCGCATTC AAGGAAGAGGAAATCGCATGATGGATCCGG AGACTACAAAGACCATGACGGTGATTATAAA GATCATGACATCGATTACAAGGATGACGATG ACAAGTCCGGATGAAAGGGCGAATTCGACC CAGCTTTCTTGTACAAAGTTGGCATTATAAG AAAGCATTGCTTATCAATTTGTTGCAACGAA CAGGTCACTATCAGTCAAAATAAAATCATTAT TTGCCATCCAGCTGATATCCCCTATAGTGAG T | gBlock to insert FLAG tag at C-terminus of pAR008_2h |

| | | |
|---|---|---|
| gBlock 1c | CTGCATTTGCCCCTCAGCTGGAGGGTAAAACGCCCGCGTACCAGGATCTGGGGCGGCCTCCCGGATCCTGGTACGCCCATGGCTGCCGACCTGGCAGCGTCCAGCACCGTGATGTGGGAACAAGATGCGGACCCCTTCGCAGGGGCAGCGGATGATTTCCCGGCATTCAACGAAGAGGAACTCGCATGGTTGATGGAGCTATTGCCTCAGGGATCCGGAGACTACAAAGACCATGACGGTGATTATAAAGATCATGACATCGATTACAAGGATGACGATGACAAGTCCGGATGAAAGGGCGAATTCGACCCAGCTTTCTTGTACAAAGTTGGCATTATAAGAAAGCATTGCTTATCAATTTGTTGCAACGAACAGGTCACTATCAGTCAAAATAAAATCATTATTTGCCATCCAGCTGATATCCCCTATAGTGAGT | gBlock to insert FLAG tag at C-terminus of pAR009_1c |

#### *Xo1 RNAseq analysis*

```
#SRR files downloaded from ENA -- repeated for all SRR files
wget ftp://ftp.sra.ebi.ac.uk/vol1/fastq/SRR932/004/SRR9320034/SRR9320034_1.fastq.gz

#Renaming files
mv SRR9320033.fastq.gz Mock3.fastq.gz
mv SRR9320034.fastq.gz 2h1.fastq.gz
mv SRR9320035.fastq.gz EV3.fastq.gz
mv SRR9320036.fastq.gz EV2.fastq.gz
mv SRR9320037.fastq.gz EV1.fastq.gz
mv SRR9320038.fastq.gz Mock2.fastq.gz
mv SRR9320039.fastq.gz Mock1.fastq.gz
mv SRR9320040.fastq.gz 2h3.fastq.gz
mv SRR9320041.fastq.gz 2h2.fastq.gz

#Nipponbare reference genome and annotation downloaded from MSU version 7.0
#genome = all.chrs.fasta
#annotation = all.gff3

#Annotation converted to .gtf and renamed
gffread all.gff3 -T -o Nippo.gtf

#The genome was indexed for STAR
STAR --runMode genomeGenerate --runThreadN 7 --genomeDir Nippo_Index --genomeFastaFiles
allchrs.fasta --sjdbGTFfile Nippo.gtf --sjdbOverhang 100

#Each pair of reads was aligned to the Nipponbare reference -- repeated for all
STAR --runThreadN 8 --runMode alignReads --genomeDir Nippo_Index/ --readFilesIn EV3_1.fastq.gz
EV3_2.fastq.gz --outSAMtype BAM SortedByCoordinate --outFileNamePrefix EV3 --readFilesCommand
zcat

#Concatenate the count columns from all STAR output to get ready for DESEQ2
paste 2h1ReadsPerGene.out.tab 2h2ReadsPerGene.out.tab 2h3ReadsPerGene.out.tab
EV1ReadsPerGene.out.tab EV2ReadsPerGene.out.tab EV3ReadsPerGene.out.tab
```

```
Mock1ReadsPerGene.out.tab Mock2ReadsPerGene.out.tab Mock3ReadsPerGene.out.tab | cut -
f1,2,6,10,14,18,22,26,30,34 | tail -n +5 > gene_count.txt

#Create a sample file to link treatment to count - it should look like this:
#Sample    Trtmnt
#2h1    2h
#2h2    2h
#2h3    2h
#EV1    EV
#EV2    EV
#EV3    EV
#Mock1    Mock
#Mock2    Mock
#Mock3    Mock

#With these files I was able to run DESEQ2 in Rstudio V1.2.5033
#note that this includes analysis such as the PCA that are not displayed in the manuscript

matrixFile <- "gene_count.txt"
sampleFile <- "samples.txt"
cts <- as.matrix(read.csv(matrixFile, sep="\t", row.names=1, header=FALSE))
coldata <- read.csv(sampleFile, sep="\t", row.names=1, header=TRUE)
#head(coldata)
#head(cts)
colnames(cts) <- rownames(coldata)

if (!requireNamespace("BiocManager", quietly = TRUE))
  install.packages("BiocManager")
if (!requireNamespace("DESeq2", quietly = TRUE))
  BiocManager::install("DESeq2")
if (!requireNamespace("dplyr", quietly = TRUE))
  install.packages("dplyr")
if (!requireNamespace("pheatmap", quietly = TRUE))
  install.packages("pheatmap")
if (!requireNamespace("tidyverse", quietly = TRUE))
  install.packages("tidyverse")

library(DESeq2)
library(dplyr)
library(pheatmap)
library(tidyverse)

dds <- DESeqDataSetFromMatrix(countData=cts, colData=coldata, design =~ Trtmnt)
dds.data <- DESeq(dds)
head(dds.data)

vsd <- vst(dds, blind=FALSE)
pca =  plotPCA(vsd, intgroup = "Trtmnt")
pca
pca$data

#Mock_vs_2h - this is Mock vs Disease
Mock.2h <- results(dds.data, contrast=c("Trtmnt", "Mock", "2h"), alpha = 0.05)
Mock.2h %>% head()
write.csv(Mock.2h, file = paste0( "Mockvs2h_results.csv"))
```

```
#Mock_vs_EV - this is Mock vs HR
Mock.EV <- results(dds.data, contrast=c("Trtmnt", "Mock", "EV"), alpha = 0.05)
Mock.EV %>% head()
write.csv(Mock.EV, file = paste0( "MockvsEV_results.csv"))

#EV_vs_2h - This is HR vs Disease
EV.2h <- results(dds.data, contrast=c("Trtmnt", "EV", "2h"), alpha = 0.05)
EV.2h %>% head()
write.csv(EV.2h, file = paste0( "EVvs2h_results.csv"))

comp = as.data.frame(Mock.2h)

hmap.query = comp %>%
  rownames_to_column("genes") %>%
  filter(., padj < 0.05 & baseMean > 1000)

vsd.table = as.data.frame(assay(vsd))
hisat2-build reconciled_assembly_v4.fa reconciled_assembly_v4
```

#### *Carolina Gold genome annotation*

```
hisat2-build reconciled_assembly_v4.fa reconciled_assembly_v4
hisat2 --max-intronlen 20000 -k 1 -p 8 --no-softclip --dta-cufflinks -x reconciled_assembly_v4 -1
Mock_1_RNAseq_forward_paired.fq.gz -2 Mock_1_RNAseq_reverse_paired.fq.gz -S
Mock_1_RNAseq.cufflinks.sam > hisat2.Mock_1.cufflinks.log 2>&1 &
hisat2 --max-intronlen 20000 -k 1 -p 8 --no-softclip --dta-cufflinks -x reconciled_assembly_v4 -1
Mock_2_RNAseq_forward_paired.fq.gz -2 Mock_2_RNAseq_reverse_paired.fq.gz -S
Mock_2_RNAseq.cufflinks.sam > hisat2.Mock_2.cufflinks.log 2>&1 &
hisat2 --max-intronlen 20000 -k 1 -p 8 --no-softclip --dta-cufflinks -x reconciled_assembly_v4 -1
Mock_3_RNAseq_forward_paired.fq.gz -2 Mock_3_RNAseq_reverse_paired.fq.gz -S
Mock_3_RNAseq.cufflinks.sam > hisat2.Mock_3.cufflinks.log 2>&1 &

samtools view -F 4 -Shub Mock_1_RNAseq.cufflinks.sam > Mock_1_RNAseq.cufflinks.bam
samtools sort -o Mock_1_RNAseq.cufflinks.sorted.bam Mock_1_RNAseq.cufflinks.bam

samtools view -F 4 -Shub Mock_2_RNAseq.cufflinks.sam > Mock_2_RNAseq.cufflinks.bam
samtools sort -o Mock_2_RNAseq.cufflinks.sorted.bam Mock_2_RNAseq.cufflinks.bam

samtools view -F 4 -Shub Mock_3_RNAseq.cufflinks.sam > Mock_3_RNAseq.cufflinks.bam
samtools sort -o Mock_3_RNAseq.cufflinks.sorted.bam Mock_3_RNAseq.cufflinks.bam

samtools merge Mock_RNAseq.cufflinks.sorted.bam Mock_1_RNAseq.cufflinks.sorted.bam
Mock_2_RNAseq.cufflinks.sorted.bam Mock_3_RNAseq.cufflinks.sorted.bam

cufflinks -p 8 Mock_RNAseq.cufflinks.sorted.bam > cufflinks.log 2>&1 &
gffread transcripts_Mock.gtf -g reconciled_assembly_v4.fa -w transcripts_Mock.fa
TransDecoder.LongOrfs -t transcripts_Mock.fa
./interproscan.sh --output-dir . --input longest_orfs.pep --iprlookup --seqtype p --appl
Coils,Gene3D,ProSitePatterns,Pfam,PANTHER,SUPERFAMILY > longest_orfs_interproscan.log 2>&1 &

#this was repeated for the 2h and EV datasets
```

#### *Heatmap and PCA generation in R*

```
#Heatmap All HR DEGs
HR_NA=read.table("Table_S1_All_HR_DEGs.txt", sep="\t", header=TRUE, row.names=1)
HR_NA_matrix=data.matrix(HR_NA)
superheat(HR_NA_matrix, heat.na.col = "black", title="All HR DEGs",
        title.size = 8, grid.vline.col = "white")

#Heatmap Only GO:0006952
Def_DEGs=read.table("Table_S2_Defense_HR_DEGs.txt", sep="\t", header=TRUE, row.names=1)
Def_DEGs_matrix=data.matrix(Def_DEGs)
superheat(Def_DEGs_matrix, heat.na.col = "black", title="Defense DEGs",
        title.size = 8, grid.vline.col = "white")

#Heatmap to compare overlapping Defense GO HR and Disease
Overlap_GODEGs=read.table("Table_S3_CGS_Disease_and_HR_DEG_Overlap.txt", sep="\t",
header=TRUE, row.names=1)
Overlap_GODEGs_matrix=data.matrix(Overlap_GODEGs)
superheat(Overlap_GODEGs_matrix, heat.na.col = "black", title="Disease DEGs CGS Overlap",
        title.size = 8, grid.vline.col = "white")

#Principal Component Analysis
data = read.table(file="rice_pathogen_RNAseq.txt", sep="\t", header=T)
pcs=prcomp(t(x),center=T,scale.=T,retx=T)

postscript(file="rice_pathogen_RNAseq_v2.ps", width=4, height=6)
plot(pcs$x[,2],pcs$x[,1], xlab="", ylab="", main="")
dev.off()
```

### Supplemental Figures

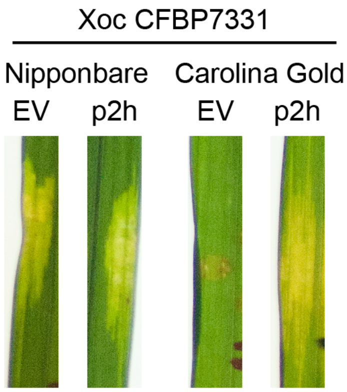


**Fig. S1.** Confirmation of CFBP7331(EV) and CFBP7331(p2h) inoculum on Nipponbare and Carolina Gold plants**.** Leaves were syringe-infiltrated with African Xoc strain CFBP7331 carrying either empty vector (EV) or *tal2h* (p2h) adjusted to $OD_{600}$ 0.4, and incubated for 10 days to allow lesion expansion. Leaves were photographed on a light box. Resistance is apparent as HR (necrosis) at the site of inoculation and disease as expanded, translucent watersoaking.

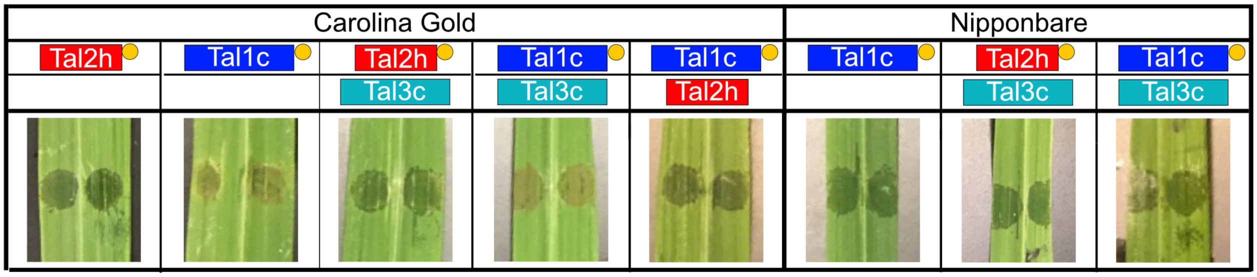


**Fig. S2.** Symptoms on Carolina Gold Select and Nipponbare leaves caused by inoculum used for the co-IP experiments. Photos were taken with overhead lighting 4 days after syringe infiltration. Resistance is apparent as HR (brown) and disease as watersoaking (dark green).

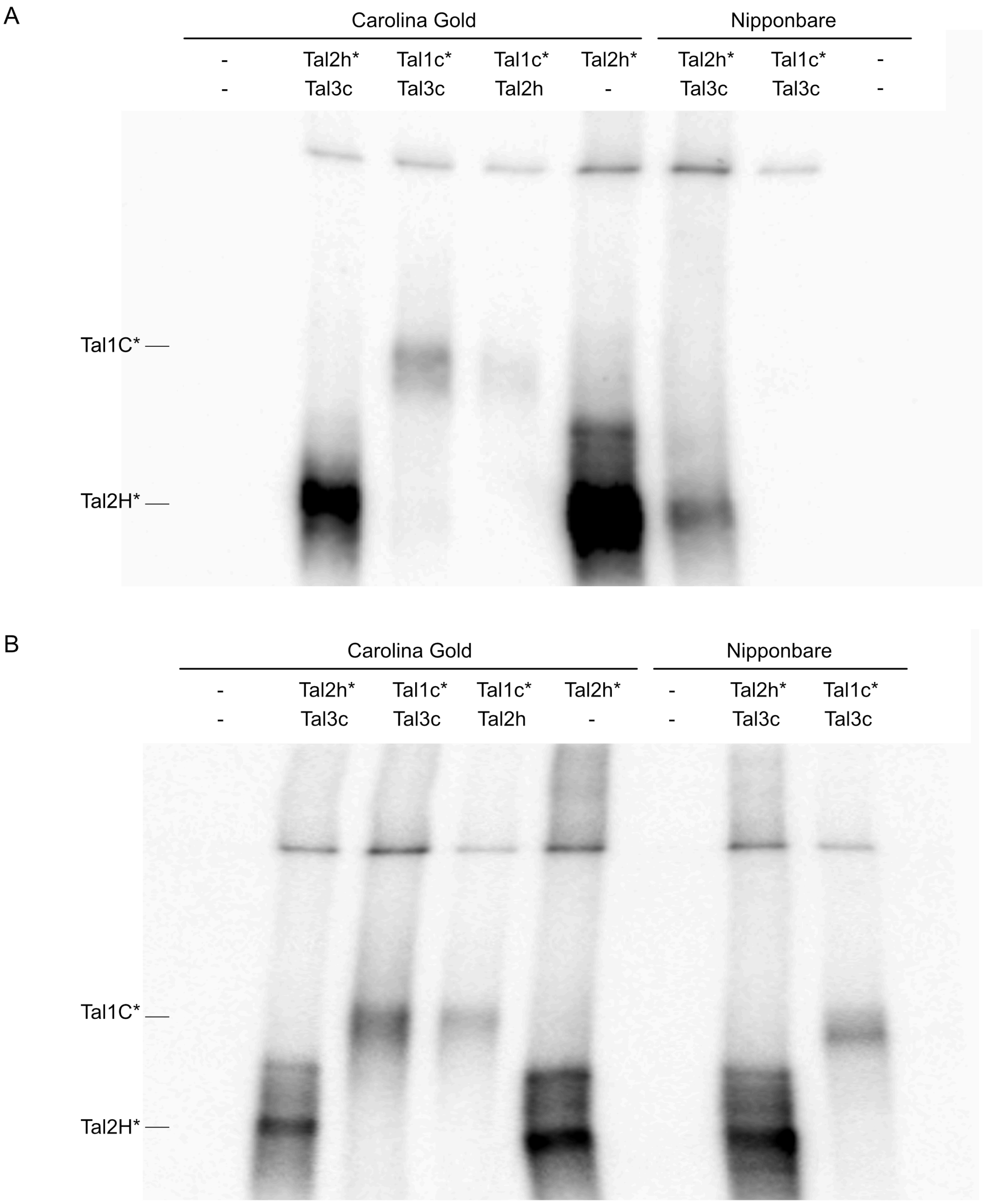


**Fig. S3.** Western blot of immunoprecipitates using anti-TALE antibody. Aliquots of the eluted immunoprecipitates from co-immunoprecipitation experiments 1 (A) and 2 (B) were resolved by 7.5% Tris-Glycine SDS-PAGE and probed with anti-TALE antibody. An asterisk indicates a 3x FLAG fusion.

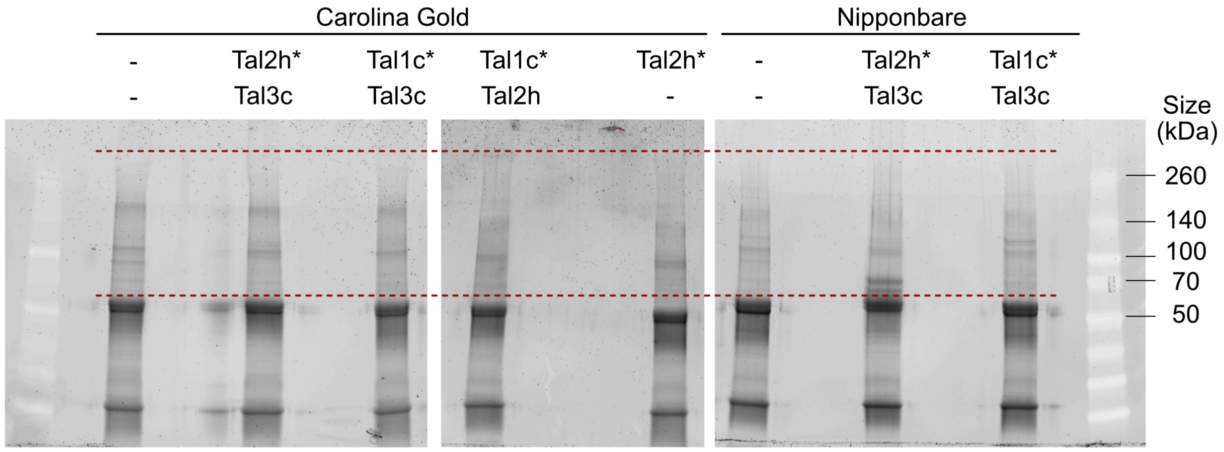


**Fig. S4.** SDS-PAGE of immunoprecipitates and size range excised for mass spectrometry.
A 4-20% polyacrylamide gel was used and proteins were stained using SYPRO Ruby. Samples were placed a lane apart to avoid cross-contamination. Red lines represent the region of the gel that were excised, containing proteins between approximately 60 and 300 kDa, and digested before mass spectrometry analysis. This gel is from the second co-immunoprecipitation experiment.

### Supplemental References


Barter, R.L., and Yu, B. 2018. Superheat: An R package for creating beautiful and extendable heatmaps for visualizing complex data. J. Comput. Graph Stat. 27:910-922.

Cermak, T., Doyle, E.L., Christian, M., Wang, L., Zhang, Y., Schmidt, C., Baller, J.A., Somia, N.V., Bogdanove, A.J., and Voytas, D.F. 2011. Efficient design and assembly of custom TALEN and other TAL effector-based constructs for DNA targeting. Nucleic Acids Res. 39:e82.

Cernadas, R.A., Doyle, E.L., Nino-Liu, D.O., Wilkins, K.E., Bancroft, T., Wang, L., Schmidt, C.L., Caldo, R., Yang, B., White, F.F., Nettleton, D., Wise, R.P., and Bogdanove, A.J. 2014. Code-assisted discovery of TAL effector targets in bacterial leaf streak of rice reveals contrast with bacterial blight and a novel susceptibility gene. PLoS Path. 10:e1003972.

Davis, S., and Meltzer, P.S. 2007. GEOquery: a bridge between the Gene Expression Omnibus (GEO) and BioConductor. Bioinformatics 23:1846-1847.

Dobin, A., Davis, C.A., Schlesinger, F., Drenkow, J., Zaleski, C., Jha, S., Batut, P., Chaisson, M., and Gingeras, T.R. 2013. STAR: ultrafast universal RNA-seq aligner. Bioinformatics 29:15-21.

Ivanov, S., and Harrison, M.J. 2014. A set of fluorescent protein-based markers expressed from constitutive and arbuscular mycorrhiza-inducible promoters to

label organelles, membranes and cytoskeletal elements in *Medicago truncatula*. Plant J. 80:1151-1163.
Karimi, M., De Meyer, B., and Hilson, P. 2005. Modular cloning in plant cells. Trends Plant Sci. 10:103-105.
Kawahara, Y., de la Bastide, M., Hamilton, J.P., Kanamori, H., McCombie, W.R., Ouyang, S., Schwartz, D.C., Tanaka, T., Wu, J., and Zhou, S. 2013. Improvement of the *Oryza sativa* Nipponbare reference genome using next generation sequence and optical map data. Rice 6:4.
Kim, D., Paggi, J.M., Park, C., Bennett, C., and Salzberg, S.L. 2019. Graph-based genome alignment and genotyping with HISAT2 and HISAT-genotype. Nat. Biotechnol. 37:907-915.
Love, M.I., Huber, W., and Anders, S. 2014. Moderated estimation of fold change and dispersion for RNA-seq data with DESeq2. Genome Biol. 15:550.
Nakagawa, T., Suzuki, T., Murata, S., Nakamura, S., Hino, T., Maeo, K., Tabata, R., Kawai, T., Tanaka, K., Niwa, Y., Watanabe, Y., Nakamura, K., Kimura, T., and Ishiguro, S. 2007. Improved Gateway binary vectors: high-performance vectors for creation of fusion constructs in transgenic analysis of plants. Biosci. Biotechnol. Biochem. 71:2095-2100.
Read, A.C., Rinaldi, F.C., Hutin, M., He, Y.-Q., Triplett, L.R., and Bogdanove, A.J. 2016. Suppression of *Xo1*-mediated disease resistance in rice by a truncated, non-DNA-binding TAL effector of *Xanthomonas oryzae*. Front. Plant Sci. 7:1516.
Read, A.C., Moscou, M.J., Zimin, A.V., Pertea, G., Meyer, R.S., Purugganan, M.D., Leach, J.E., Triplett, L.R., Salzberg, S.L., and Bogdanove, A.J. 2020. Genome assembly and characterization of a complex zfBED-NLR gene-containing disease resistance locus in Carolina Gold Select rice with Nanopore sequencing. PLoS Genet. 16:e1008571.
Reimers, P.J., Guo, A., and Leach, J.E. 1992. Increased activity of a cationic peroxidase associated with an incompatible interaction between *Xanthomonas oryzae* pv. oryzae and rice *Oryza sativa*. Plant Physiol. 99:1044-1050.
Rinaldi, F.C., Doyle, L.A., Stoddard, B.L., and Bogdanove, A.J. 2017. The effect of increasing numbers of repeats on TAL effector DNA binding specificity. Nucleic Acids Res. 45:6960-6970.
Schindelin, J., Arganda-Carreras, I., Frise, E., Kaynig, V., Longair, M., Pietzsch, T., Preibisch, S., Rueden, C., Saalfeld, S., Schmid, B., Tinevez, J.Y., White, D.J., Hartenstein, V., Eliceiri, K., Tomancak, P., and Cardona, A. 2012. Fiji: an open-source platform for biological-image analysis. Nat. Methods 9:676-682.
Smyth, G.K. 2004. Linear models and empirical bayes methods for assessing differential expression in microarray experiments. Stat. Appl. Genet. Mol. Biol. 3:Article3.
Tanabe, S., Yokotani, N., Nagata, T., Fujisawa, Y., Jiang, C., Abe, K., Ichikawa, H., Mitsuda, N., Ohme-Takagi, M., Nishizawa, Y., and Minami, E. 2014. Spatial regulation of defense-related genes revealed by expression analysis using dissected tissues of rice leaves inoculated with *Magnaporthe oryzae*. J. Plant Physiol. Pathol. 2:1000135.
Tanaka, Y., Nakamura, S., Kawamukai, M., Koizumi, N., and Nakagawa, T. 2011. Development of a series of gateway binary vectors possessing a tunicamycin

resistance gene as a marker for the transformation of *Arabidopsis thaliana*. Biosci. Biotechnol. Biochem. 75:804-807.

Tariq, R., Wang, C., Qin, T., Xu, F., Tang, Y., Gao, Y., Ji, Z., and Zhao, K. 2018. Comparative transcriptome profiling of rice near-isogenic line carrying *Xa23* under infection of *Xanthomonas oryzae* pv. oryzae. Int. J. Mol. Sci. 19:717.

Thomas, C.J., Cleland, T.P., Zhang, S., Gundberg, C.M., and Vashishth, D. 2017. Identification and characterization of glycation adducts on osteocalcin. Anal. Biochem. 525:46-53.

Trapnell, C., Roberts, A., Goff, L., Pertea, G., Kim, D., Kelley, D.R., Pimentel, H., Salzberg, S.L., Rinn, J.L., and Pachter, L. 2012. Differential gene and transcript expression analysis of RNA-seq experiments with TopHat and Cufflinks. Nat. Protoc. 7:562-578.

Triplett, L.R., Hamilton, J.P., Buell, C.R., Tisserat, N.A., Verdier, V., Zink, F., and Leach, J.E. 2011. Genomic analysis of *Xanthomonas oryzae* isolates from rice grown in the United States reveals substantial divergence from known *X. oryzae* pathovars. Appl. Environ. Microbiol. 77:3930-3937.

Yang, Y., Anderson, E., and Zhang, S. 2018. Evaluation of six sample preparation procedures for qualitative and quantitative proteomics analysis of milk fat globule membrane. Electrophoresis 39:2332-2339.

Yang, Y., Thannhauser, T.W., Li, L., and Zhang, S. 2007. Development of an integrated approach for evaluation of 2-D gel image analysis: impact of multiple proteins in single spots on comparative proteomics in conventional 2-D gel/MALDI workflow. Electrophoresis 28:2080-2094.

Zhou, Y.L., Xu, M.R., Zhao, M.F., Xie, X.W., Zhu, L.H., Fu, B.Y., and Li, Z.K. 2010. Genome-wide gene responses in a transgenic rice line carrying the maize resistance gene *Rxo1* to the rice bacterial streak pathogen, *Xanthomonas oryzae* pv. oryzicola. BMC Genomics 11:78.
